# Split personality of Aluminum Activated Malate Transporter family proteins: facilitation of both GABA and malate transport

**DOI:** 10.1101/215335

**Authors:** Sunita A. Ramesh, Muhammad Kamran, Wendy Sullivan, Larissa Chirkova, Mamoru Okamoto, Fien Degryse, Michael McLaughlin, Matthew Gilliham, Stephen D. Tyerman

**Author notes:** The author responsible for distribution of materials integral to the findings presented in this article in accordance with the policy described in the Instructions for Authors (www.plantcell.org) is: Stephen D Tyerman.

## Abstract

Plant aluminum activated malate transporters (ALMTs) are currently classified as anion channels; they are also known to be regulated by diverse signals leading to a range of physiological responses. Gamma-aminobutyric acid (GABA) regulation of anion flux through ALMT proteins requires the presence of a specific amino acid motif in ALMTs that shares similarity with a GABA-binding site in mammalian GABA_A_ receptors. Here, we explore why TaALMT1-activation leads to a negative correlation between malate efflux and endogenous GABA concentrations ([GABA]_i_) in both wheat root tips and in heterologous expression systems. We show that TaALMT1 activation reduces [GABA]_i_ because TaALMT1 facilitates GABA efflux. TaALMT1-expression also leads to GABA transport into cells, demonstrated by a yeast complementation assay and via ^14C^GABA uptake into TaALMT1-expressing *Xenopus laevis* oocytes; this was found to be a general feature of all ALMTs we examined. Mutation of the GABA motif (TaALMT1^F213C^) prevented both GABA influx and efflux, and uncoupled the relationship between malate efflux and [GABA]_i_. We conclude that ALMTs are likely to act as both GABA and anion transporters *in planta*. GABA and malate appear to interact with ALMTs in a complex manner regulating each other’s transport, suggestive of a role for ALMTs in communicating metabolic status.

## INTRODUCTION

Gamma-aminobutyric acid (GABA) is a four carbon non-proteinogenic amino acid that was first discovered in potato tubers (Steward et al., 1949); however, since this time it has mainly been studied in mammals as an inhibitory neurotransmitter (Sigel and Steinmann, 2012). When plants encounter stress, both abiotic (e.g. hypoxia, heat, cold, salt, drought) and biotic (e.g. herbivory, pathogen infection), they rapidly accumulate GABA (Shelp et al., 2012). This accumulation has been shown to play an important role in the regulation of: C:N balance (Fait et al., 2008; Fait et al., 2011); cytosolic pH (Carroll et al., 1994; Shelp et al., 1999); salt tolerance (Renault et al., 2010); and oxidative stress tolerance (Bouche et al., 2003b; Bouche and Fromm, 2004). GABA has also been proposed to be an endogenous plant-signaling molecule (Kinnersley and Turano, 2000; Palanivelu et al., 2003; Roberts, 2007). More recently, it was shown that GABA at low micromolar concentrations regulates anion currents through <u>A</u>luminum <u>A</u>ctivated <u>M</u>alate <u>T</u>ransporters (ALMT) proteins from various species. While TaALMT1 can be activated by aluminum (Al^3+^) this is not a general feature of ALMTs; some ALMTs can facilitate anion efflux when trans-activated by external anions such as SO_4_^2−^ or malate^2−^ in alkaline solutions (Ramesh et al., 2015). A putative GABA binding motif was discovered in ALMTs with homology to the one found in mammalian GABA_A_ receptors, and treatment with micromolar concentrations of muscimol (an analogue of GABA) resulted in inhibition of anion flux (Ramesh et al., 2015). Addition of bicuculline (a GABA receptor antagonist) attenuated the effect of both muscimol and GABA (Ramesh et al., 2015). These results suggest that there are distinct classes of anion channels from plant and animal cells that have comparable modes of GABA regulation (Žárský, 2015; Gilliham and Tyerman, 2016; Ramesh et al., 2016).

It is well established that in acidic soils TaALMT1 confers Al^3+^ tolerance in wheat through exuding malate from the root tips and chelating toxic Al^3+^ (Delhaize and Ryan, 1995; Ma et al., 2001; Sasaki T, 2004). Exogenous application of GABA or muscimol to the roots of wheat seedlings with high TaALMT1 expression inhibited malate efflux and impaired root growth in the presence of Al^3+^, which phenocopied a near isogenic line with less expression of TaALMT1 and less Al^3+^ tolerance (Ramesh et al., 2015). Interestingly, in these conditions, it was observed that when root efflux of malate was high, endogenous GABA concentrations ([GABA]_i_) in the cells were low and *vice versa* (Ramesh et al., 2015). This reciprocal relationship remained unexplained and may indicate either TaALMT1 activation caused changes in [GABA]_i_ or that [GABA]_i_ is altered in some way that then regulates TaALMT1.

Identification of a putative GABA binding motif in ALMTs provided a possible mechanism by which plant GABA may act as a signal (Ramesh et al., 2015; Žárský, 2015), but we are yet to fully understand the molecular and physiological basis of how this occurs and the relationship between anion flux and GABA regulation in plant cells. A number of pharmacological agents have been used to characterise animal GABA receptors, some of which are plant or fungal derived, either as agonists i.e muscimol, or as regulators of GABA synthesis or catabolism i.e. amino-oxyacetate (AOA) and vigabatrin (Wood and Peesker, 1973; Jackson et al., 1982; Grant and Heel, 1991). Manipulating [GABA]_i_ in plant cells and studying the effect this has on anion efflux mediated by ALMTs will increase our understanding of the role of GABA in plants under different stresses.

It was previously hypothesised that ALMT might sense and signal metabolic status through their regulation by GABA and malate, which alters membrane voltage and transduces the signal into a physiological response (Gilliham and Tyerman, 2016; Xu et al., 2016). Cellular efflux of GABA has been well documented (Bown and Shelp, 1989; Chung et al., 1992). Micromolar concentrations of GABA are found in root exudates and the apoplast, and among all amino acids exuded from wheat roots, GABA shows the highest efflux (Warren, 2015); it was envisaged that such carbon and nitrogen loss might only be justified energetically if GABA was involved in signaling (Gilliham and Tyerman, 2016). While a high affinity GABA influx transporter (AtGAT1) has been characterised and is expressed in Arabidopsis roots (Meyer et al., 2006), no transporter that can efflux GABA from the cytoplasm into the apoplast has been identified. This raises an interesting question as to how GABA exits the cytoplasm and enters the apoplast.

In this study we used two GABA analogs to manipulate [GABA]_i_ in cells expressing TaALMT1: vigabatrin – a GABA transaminase (GABA-T) inhibitor used as an antiepileptic in humans (Livingston et al., 1989; Nanavati and Silverman, 1991), and amino-oxyacetate (AOA) – an inhibitor of both GABA-T and glutamate decarboxylase (GAD) (John and Charteris, 1978; Miller et al., 1991; Snedden et al., 1992). We demonstrate that exogenous application of AOA and vigabatrin has an affect upon anion efflux via TaALMT1. This led to negative correlations between [GABA]_i_ and anion efflux being observed, which are also evident following Al^3+^ application. Interestingly, these negative correlations are uncoupled when the site directed mutant TaALMT1^F213C^ – impaired in its GABA regulation of malate transport – is expressed instead of TaALMT1. Further, we propose that the reduction in [GABA]_i_ upon Al^3+^ treatment at low pH is likely to be due, in part, to efflux of GABA via TaALMT1. More broadly, our work reveals that changes in intracellular and apoplasmic concentrations of GABA can be facilitated by an ALMT protein, resulting in exquisite regulation of malate efflux via both a feed-forward (malate) and feed-back (GABA) regulation. Such ALMT activity will strongly affect [GABA]_i_, resulting in changes in metabolic flux through the GABA shunt. In the case of the wheat root this provides a likely signalling mechanism by which TaALMT1 can regulate root growth in acidic and alkaline conditions.

## RESULTS

### Validating measurement of intracellular GABA concentration

We routinetly use a GABase enzyme assay to measure GABA concentration, which utilises a GABA-T enzyme. Whilst examining the effects of inhibitors on cell and tissue GABA concentrations ([GABA]_i_) we found that this assay was inhibited by AOA when added directly to the enzyme mix. We therefore examined if the GABase assay was compromised when it was used on extracts of tissues that had been treated with AOA, Al^3+^ or vigabatrin by performing GABA spike and recovery experiments (Supplemental Figure 1). After 22 h treatment with 1 mM AOA the recovery of spiked GABA from wheat seedling root extract was 97% (Supplemental Figure 1A,B,C); Al^3+^ treatment also did not compromise measurement of [GABA]_i_ (Supplemental Figure 1D,E,F). Vigabatrin treated wheat root tips also yielded full recovery of spiked GABA (Supplemental Figure 1G,H,I). *Xenopus laevis* oocytes treated with these compounds were also examined and similarly full recovery of GABA was obtained (Supplemental Figure 1J,K,L). We also used Ultra High Performance Liquid Chromatography (UPLC) to measure [GABA]_i_ on the same wheat root tip tissue extracts, after treatment with Al^3+^ and AOA, validating the results obtained by the enzyme assay (Supplemental Figure 1M). Thus, we expect the GABase assay to faithfully measure [GABA]_i_ following Al^3+^, AOA or vigabatrin treatments.

### Wheat roots malate efflux and [GABA]_i_

External Al^3+^ at low pH was previously shown to reduce [GABA]_i_ while stimulating malate efflux from roots of wheat seedlings (Ramesh et al. 2015). The Al^3+^ tolerant wheat line ET8 has higher expression of *TaALMT1* that is increase by external Al^3+^ compared to ES8, its near isogenic line (NIL) (Yamaguchi et al., 2005) (Supplemental Figure 2). We confirmed that ET8 exhibited Al^3+^ (100 µM) induced malate efflux from roots of 3-day old wheat seedlings and that this co-incided with a reduction in root tip [GABA]_i_ (Figure 1). Line ES8 had lower [GABA]_i_ than ET8 and ES8 [GABA]_i_ did not respond to Al^3+^ treatment (Supplementary Figure 1M). When corresponding values from the same plants were plotted against each other we observed that Al^3+^-stimulated malate efflux from roots had a negative linear relationship with root tip [GABA]_i_ (Figure 1A). Root tips have previously been shown to be the major site of malate efflux in response to Al^3+^ (Delhaize et al., 1993). We confirmed this response (Figure 1B), which also corresponded to reduced root tip [GABA]_i_ (Figure 1C). Amino-oxyacetate (AOA) treatment (1 mM) at pH 4.5 also stimulated malate efflux (Figure 1B) and reduced [GABA]_i_ in excised root tips of ET8 seedlings (Figure 1C) to the same degree as 100 µM Al^3+^.

**Figure 1.**
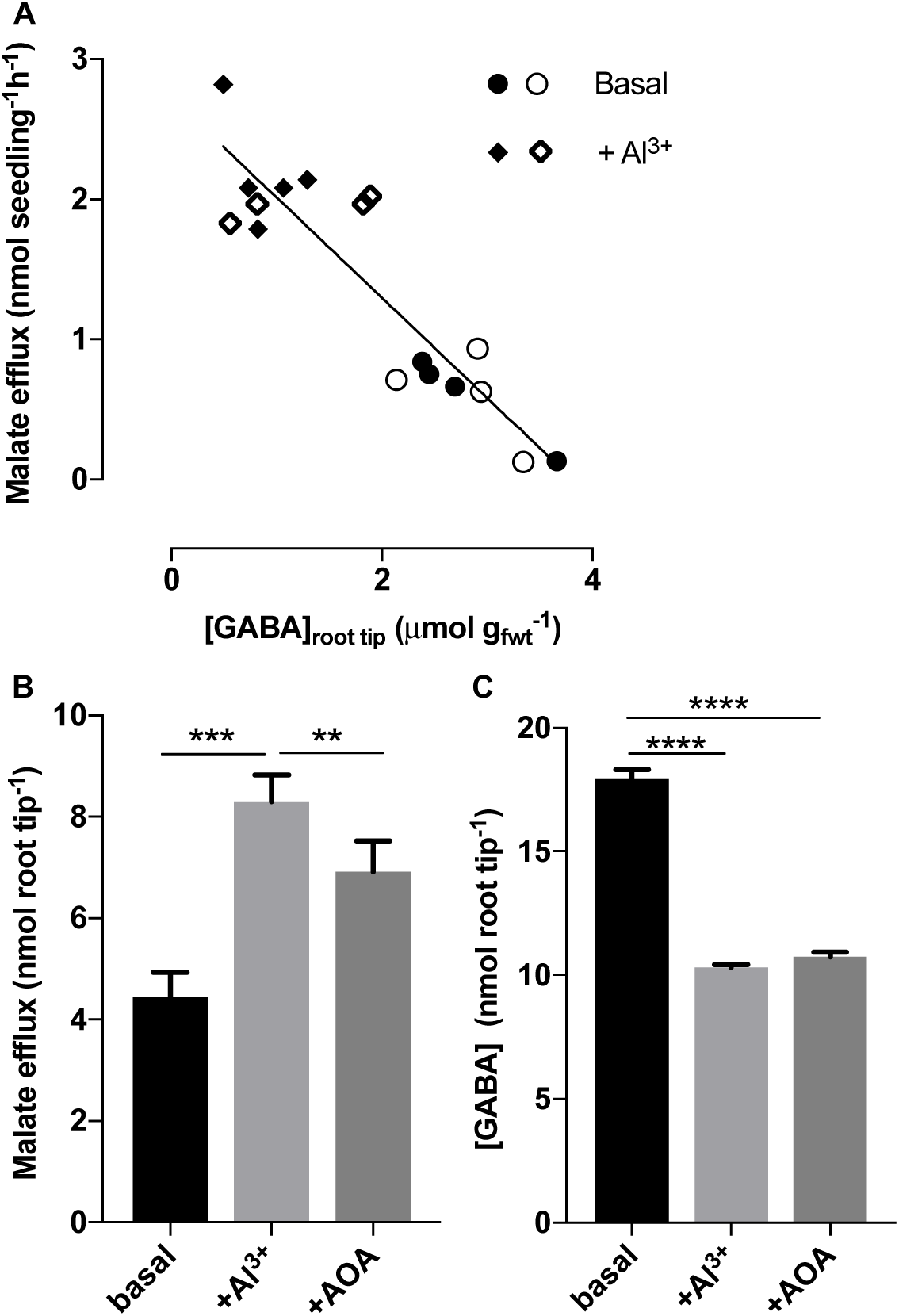
Aluminum and aminooxyacetate (AOA) stimulate efflux of malate from roots and root tips of ET8 wheat seedlings at pH 4.5 and decrease [GABA]_i_ in root tips. **(A)** Malate efflux over 22 h from intact seedling root tips correlates with [GABA]_i_ in root tips of ET8 in response to Al^3+^ at pH 4.5. Linear regression (Y = −0.7147*X + 2.727), R^2^ = 0.82, P < 0.0001. **(B)** Malate efflux from excised root tips of wheat seedlings (NIL line ET8) after 6 h in basal solution (pH 4.5), and addition of Al^3+^ (100 μM) or AOA (1 mM). **(C)** [GABA]_i_ in excised root tips in response to the same treatments. A n=8-9, B & C n=10, replicates from 2 experiments and *, **, *** indicate significant differences between treatments at P< 0.05, 0.01, 0.001 respectively, using a one-way ANOVA.

We treated roots of intact ET8 wheat seedlings with either a basal solution at pH 4.5 or pH 7.5 ± 10 mM Na_2_SO_4_ (to add a trans-activating anion) ± 1 mM AOA (at pH 4.5) or ± 100 µM vigabatrin (at pH 7.5) (Figure 2). Vigabatrin was only used at pH 7.5 since according to its mode of action it should increase [GABA]_i_, which would be expected to inhibit a transactivated malate efflux via TaALMT1. Both in the presence and absence of external SO_4_^2−^, AOA significantly stimulated malate efflux at pH 4.5 (Figure 2A) – this confirmed the results we observed for excised root tips (Figure 1). The corresponding [GABA]_i_ of root tips were lower in the presence of AOA (Figure 2B) appearing to confirm a mode of action of AOA as a GAD inhibitor. The presence of SO_4_^2−^ at pH 4.5 made no difference to root tip [GABA]_i_. Taking all the individual replicates together for two independent experiments a significant negative correlation was observed between malate efflux and root tip [GABA]_i_ at pH 4.5 (Figure 2C). At pH 7.5, malate efflux was significantly higher in the presence of SO_4_^2−^ but vigabatrin inhibited the SO_4_^2−^ stimulated efflux (Figure 2D). Root tip [GABA]_i_ at pH 7.5 was slightly lower but not significantly so than at pH 4.5 (Figure 2B and E), but was significantly decreased by the addition of SO_4_^2−^ contrasting with the lack of effect seen at pH 4.5. Treatment with SO_4_^2−^ plus vigabatrin significantly increased [GABA]_i_ from that found following SO_4_^2−^ treatment alone (Figure 2E), consistent with its proposed action of inhibiting GABA catabolism via GABA-T. Again there was a significant negative linear relationship between root tip [GABA]_i_ and malate efflux (Figure 2F), though the slope was lower than that observed at pH 4.5 (P<0.001). Taken together, these results show that in roots of intact wheat seedlings there exists a strong and consistent correlation between malate flux and [GABA]_i_ in root tips both at low and high pH regardless of the treatment that affects malate efflux.

**Figure 2.**
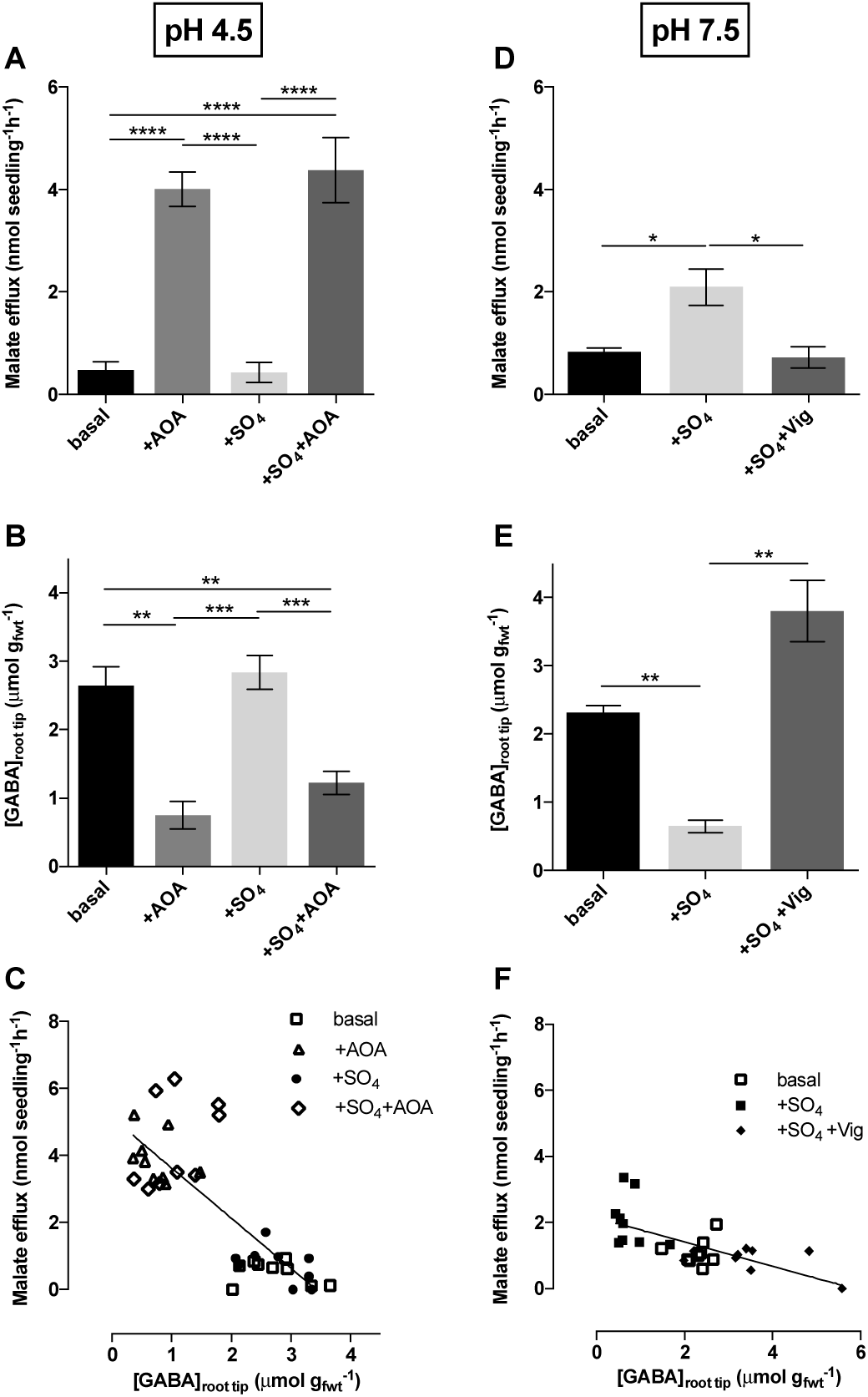
Malate efflux activation and inhibition at low and high pH by AOA and vigabatrin is coupled to root tip [GABA]_i_ in intact roots of ET8 wheat seedlings. **(A)** Malate efflux at pH 4.5 in basal solution plus AOA (1 mM), Na_2_SO_4_ (10 mM) and AOA + Na_2_SO_4_. **(B)** Root tip [GABA]_i_ measured at the end of the flux period (22h) with the same treatments above. **(C)** Negative linear correlation between malate efflux and root tip [GABA]_i_ for the individual replicates for the treatments at pH 4.5. Linear regression (Y = −1.492*X + 5.12), R^2^ = 0.64, P < 0.0001 **(D)** Malate efflux at pH 7.5 in basal solution plus Na_2_SO_4_ (10 mM) and Vigabatrin (100 µM) + Na_2_SO_4_. **(E)** Root tip [GABA]_i_ measured at the end of the flux period (22h) with the same treatments above. **(F)** Negative linear correlation between malate efflux and root tip [GABA]_i_ for the individual replicates for the treatments at pH 7.5. Linear regression (Y = −0.3662*X + 2.138), R^2^ = 0.44, P = 0.0002. *, **, ***, **** indicate significant differences between treatments at P< 0.05, 0.01, 0.001, 0.0001 respectively, using a one-way ANOVA.

### Malate efflux and [GABA]_i_ in tobacco BY2 expressing *TaALMT1*

Tobacco BY2 cells were used to see if heterologous expression of *TaALMT1* was sufficient to induce the negative linear relationship between malate efflux and [GABA]_i_ that was observed in wheat roots. AOA greatly stimulated malate efflux in *TaALMT1*-expressing BY2 cells (Figure 3A, Supplemental Figure 3C); the effect being greatest at pH 4.5 compared to higher pHs (Supplemental Figure 4). At pH 4.5, the stimulated malate efflux corresponded to a lowered [GABA]_i_ at the end of the efflux period (Figure 3B). There was no effect of SO_4_^2−^ by itself on either parameter at pH 4.5 (Figure 3A,B). Both malate efflux and [GABA]_i_ did not vary in BY2 cells transformed with the empty vector when treated with AOA or Al^3+^ (Supplemental Figure 3). As observed for wheat roots, BY2 cells expressing *TaALMT1* exhibited a significant negative linear correlation between malate efflux and [GABA]_i_ across individual replicates and treatments at pH 4.5 (Figure 3C, Supplemental Figure 5).

**Figure 3.**
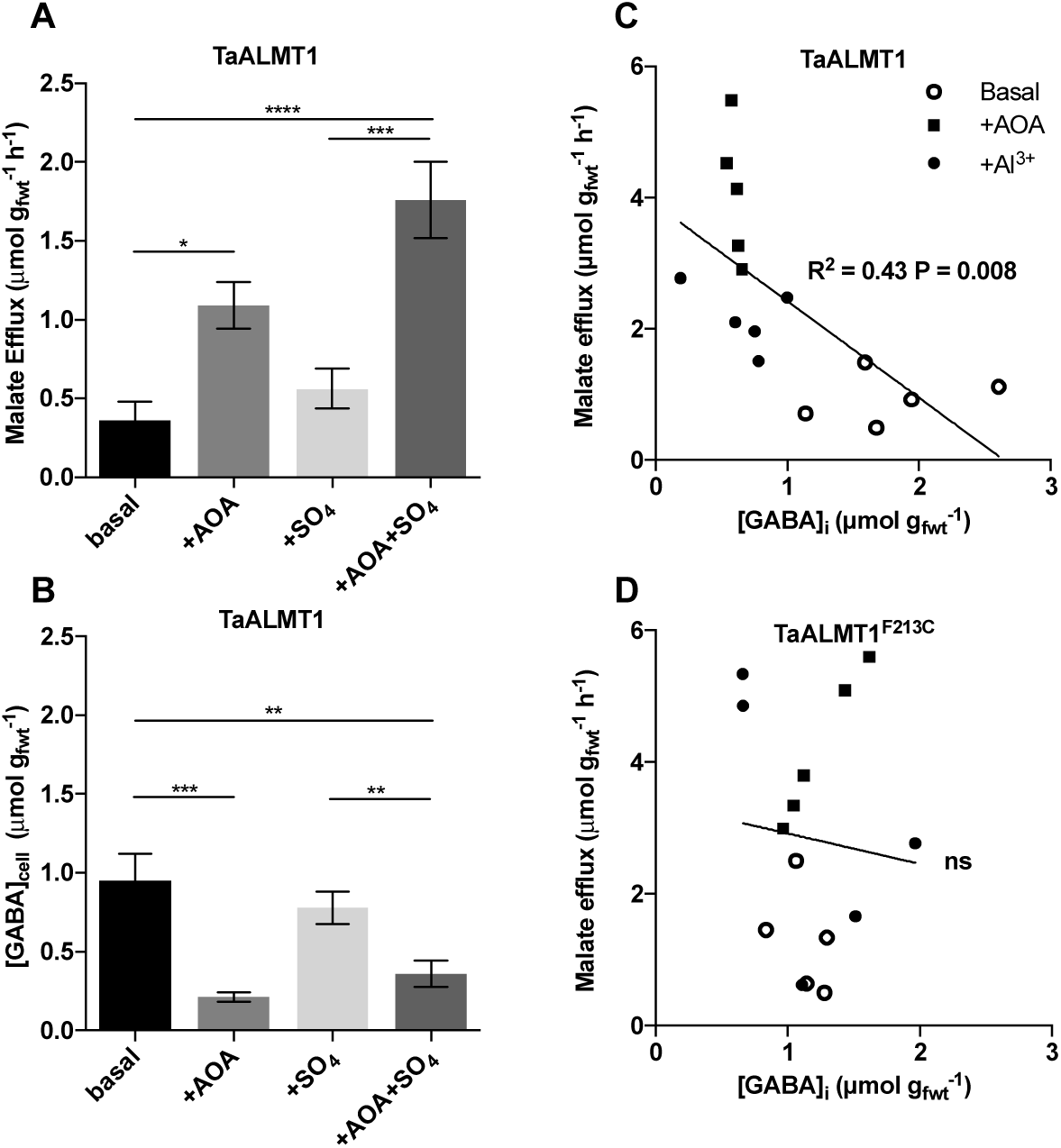
Tobacco BY2 cells expressing TaALMT1 but not TaALMT1^F213C^ show the same reciprocal response between malate efflux and endogenous GABA concentration in response to AOA and Al^3+^ at pH 4.5 as intact ET8 roots. **(A)** AOA (1 mM) activates malate efflux from BY2 cells expressing TaALMT1 at pH 4.5 in standard (basal) solution over 22 h. There was no effect of AOA on malate efflux from BY2 cells expressing empty vector (Supplemental Figure 4A). **(B)** [GABA]_i_ in BY2 cells expressing TaALMT1 at the end of the efflux period. There was no effect of AOA on [GABA]_i_ in BY2 cells expressing empty vector (Supplemental Figure 3D,E,F). All data *n*=5 replicates; *, **, ***, **** indicate significant differences between treatments at P< 0.05, 0.01, 0.001, 0.0001 respectively, using a one-way ANOVA. **(C)** WT TaALMT1 malate efflux versus [GABA]_i_ in the cells at the end of the efflux period (22 h) at pH 4.5 after treatment with 100 µM Al^3+^ or 1 mM AOA addition to basal solution (different experiment to A and B). Linear regression (Y = −1.471*X + 3.898), R^2^ = 0.43, P = 0.008. **(D)** Site directed mutant TaALMT1^F213C^ at pH 4.5, treatments as in (C); ns=regression not significant.

In contrast to pH 4.5, SO_4_^2−^ at pH 7.5 significantly stimulated malate efflux from BY2 cells expressing *TaALMT1* (Supplemental Figure 6A). The addition of vigabatrin inhibited SO_4_^2−^ stimulated malate efflux in a dose dependent manner with complete block observed at 100 µM (Supplemental Figure 6A), but had no effect on empty vector expressing controls (Supplemental Figure 6C). In BY2 cells expressing *TaALMT1* the reduction of malate efflux correlated with an increase in [GABA]_i_ (Supplemental Figure 6B). [GABA]_i_ in BY2 cells were elevated by vigabatrin as expected from its proposed action on GABA-T in both *TaALMT1* (Supplemental Figure 6B) and empty vector expressing cells (Supplemental Figure 6D). Therefore, following both AOA and vigabatrin treatment the significant relationships between malate efflux and [GABA]_i_ that were observed were dependent upon the expression of *TaALMT1*.

### The F213C mutation in TaALMT1 uncouples the relationship between [GABA]_i_ and malate efflux

A phenylalanine residue (F) has been shown to be important for GABA sensitivity in GABA_A_ receptors and TaALMT1 (Boileau et al., 1999; Ramesh et al., 2015). Mutation of this residue abolishes GABA sensitivity in animals and impairs GABA sensitivity of TaALMT1 but did not abolish activation of the malate efflux by Al^3+^ or external anions (Ramesh et al., 2015). Therefore, we tested whether the exogenous application of Al^3+^ or AOA affected the relationship between malate efflux and [GABA]_i_ in tobacco BY2 cells expressing *TaALMT1* and the site directed mutant *TaALMT1^F213C^*. In the presence of Al^3+^ or AOA at low pH, malate efflux and [GABA]_i_ in the *TaALMT1* expressing cells were significantly negatively correlated (Figure 3C) but these were uncoupled in cells expressing the *TaALMT1*^*F213C*^ mutant at pH 4.5 (Figure 3D). Malate efflux increased with exogenous application of Al^3+^ or AOA in cells expressing *TaALMT1*^*F213C*^ similarly to cells expressing *TaALMT1* (Supplemental Figure 3A,B,C), but unlike for *TaALMT1* expressing cells, [GABA]_i_ was not significantly reduced by these treatments (Supplemental Figure 3D, E,F). At pH 7.5, malate efflux from *TaALMT1*^*F213C*^ and *TaALMT1* expressing cells was stimulated by SO_4_^2−^ to a similar degree (Supplemental Figure 7A,B,C), but, in contrast to cells expressing *TaALMT1,* vigabatrin did not inhibit efflux from cells expressing *TaALMT1*^*F213C*^ even though [GABA]_i_ was elevated in both (Supplemental Figure 7D,E,F). Thus, the relationship between malate efflux and [GABA]_i_ was also uncoupled in cells expressing the *TaALMT1*^*F213C*^ mutant at pH 7.5 (Supplemental Figure 7G,H). The empty vector controls also showed increased [GABA]_i_ with vigabatrin treatment (Supplemental Figure 7D,E,F) but did not show a significant correlation between malate efflux and [GABA]_i_ (Supplemental Figure 8).

### Activation of TaALMT1 results in GABA efflux

To determine if the reduction in [GABA]_i_ was due to transport of GABA through TaALMT1, we tested if Al^3+^ not only activated malate efflux but also GABA efflux from the wheat NILs ET8 and ES8 (Figure 4). We observed that ET8 showed not only significantly higher malate efflux (Figure 4A) but also higher GABA efflux when compared to ES8 (Figure 4B). Similarly transgenic barley expressing *TaALMT1* (OE) (Delhaize et al., 2004; Ramesh et al., 2015) when exposed to Al^3+^ at low pH, showed significantly increased malate efflux (Figure 4C) as well as higher GABA efflux (Figure 4D) when compared to the Golden Promise background alone. It is also interesting to note that the efflux of GABA was higher than that of malate on a molar basis by 2-fold in ET8 wheat and over 500-fold in *TaALMT1* expressing barley roots. GABA efflux was also examined in tobacco BY2 cells expressing *TaALMT1* and the *TaALMT1*^*F213C*^ mutant where at low pH we observed very large GABA efflux with Al^3+^ treatment only for cells expressing *TaALMT1* (Figure 4E). GABA efflux in response to Al^3+^ from BY2 cells expressing *TaALMT1* was also higher than malate efflux (compare Figure 4E with Figure 3C or Supplemental Figure 3B).

**Figure 4.**
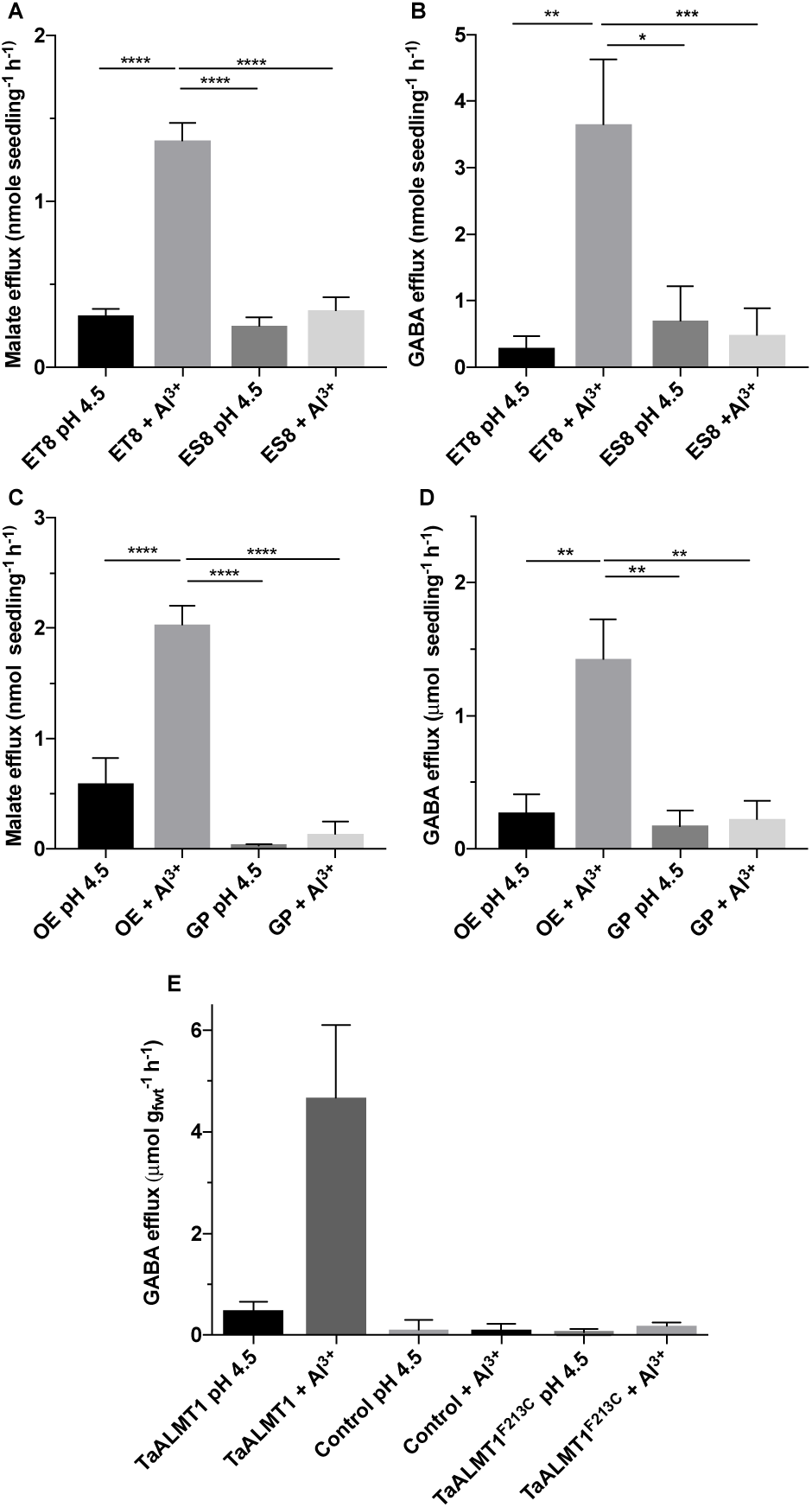
A large GABA efflux occurs from roots and BY2 cells expressing TaALMT1 together with malate efflux in response to Al^3+^ at pH 4.5. **(A)** Increased malate efflux is observed from intact seedling roots of NIL ET8 but not ES8 wheat after treatment with 100 µM Al^3+^. **(B)** Increased GABA efflux is observed from roots of ET8 but not ES8 in response to Al^3+^. **(C)** Increased malate efflux is observed from intact seedling roots of transgenic barley (Golden Promise) expressing TaALMT1 (OE) compared with Golden Promise (GP) background after treatment with 100 µM Al^3+^. **(D)** Increased GABA efflux is observed from barley roots TaALMT1 OE but not GP as above. **(E)** GABA efflux from tobacco BY2 cells over 22 h in response to Al^3+^ at pH 4.5 for empty vector (Control), cells expressing TaALMT1 or site directed mutant TaALMT1^F213C^. Only TaALMT1 + Al^3+^ showed a significant increase in GABA efflux (P<0.0001). For malate efflux see Supplemental Figure 3. All data *n*=5-12 replicates; *, **, ***, **** indicate significant differences between treatments at P< 0.05, 0.01, 0.001, 0.0001 respectively, using a one-way ANOVA.

Given that Al^3+^ activates GABA efflux through TaALMT1 in addition to malate, we tested the possibility that GABA may also complex Al^3+^, since there was no data in the literature on possible interactions between GABA and Al^3+^. Using a fluoride competitive ligand method and comparing GABA with the organic anions citrate, oxalate, malate and salicylate, we found that GABA had very low affinity for Al^3+^ compared to these other compounds, with the strength of complexation following the order: citrate>oxalate>malate>salicylate>>GABA (Detailed in Supplemental material A). We also found no synergistic or antagonistic interaction between GABA and malate in Al^3+^ complexation.

### AOA and vigabatrin have direct effects on TaALMT1

Since AOA had similar effects to Al^3+^ in activation of malate efflux and on depression [GABA]_i_, it prompted us to examine if the GABA analogs AOA and vigabatrin may have direct effects on TaALMT1 that may then alter [GABA]_I_ (Figure 5). Using TEVC on *X. laevis* oocytes expressing TaALMT1 and TaALMT1^F213C^, we perfused the bath with 1 mM AOA and 100 µM Al^3+^ to compare the inward current activation corresponding to activation of malate efflux (Figure 5A,B). AOA activates TaALMT1 and the TaALMT1^F213C^ mutant rapidly and with similar kinetics to Al^3+^. Vigabatrin was also examined for its effect on anion-activated currents at pH 7.5 (Figure 5C). Vigabatrin also acted rapidly on the malate-activated inward current giving 100% inhibition at the applied concentration of 100 µM. Furthermore, this inhibition was consistent for repeated applications indicating no carry over effect that would be expected if [GABA]_i_ were increased by inhibition of GABA transaminase. Oocyte [GABA]_i_ were examined in separate batches of oocytes after 10 mins treatment with Al^3+^, AOA and vigabatrin and compared to water injected controls (Figure 5E,F,G). Both Al^3+^ and AOA resulted in reduced [GABA]_i_, but only in *TaALMT1* expressing oocytes. The effect of vigabatrin was examined at pH 7.5. In this case we had previously shown that Na_2_SO_4_ addition stimulated malate currents (Ramesh et al., 2015). Vigabatrin increased [GABA]_i_ compared to Na_2_SO_4_ treated oocytes only in *TaALMT1* expressing oocytes. Taken together, these data are consistent with rapid effects of both AOA and vigabatrin on TaALMT1, which appears to alter [GABA]_i_ by either activation of GABA efflux through TaALMT1 (AOA) or inhibition (vigabatrin).

**Figure 5.**
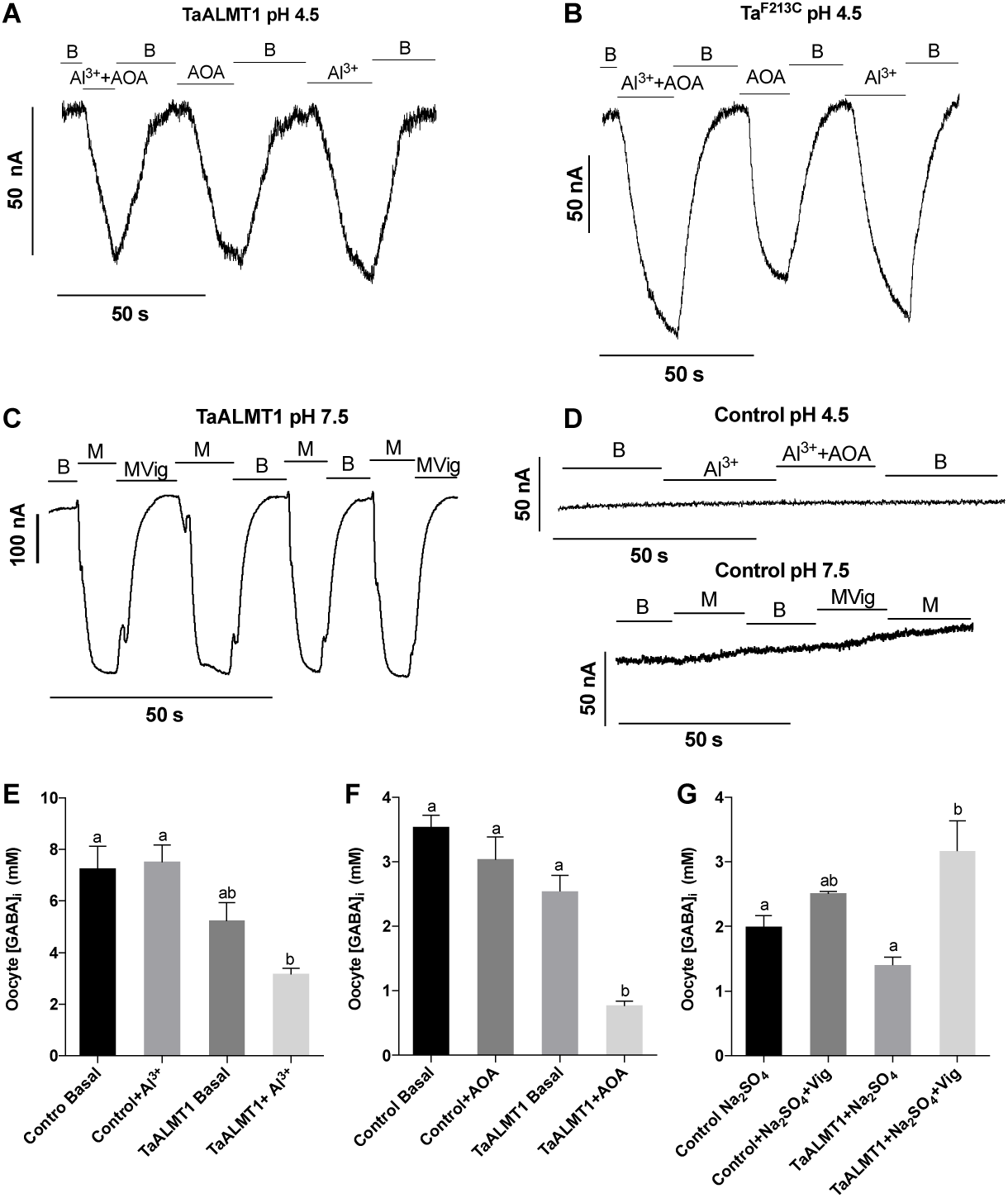
Rapid activation of inward currents (anion efflux) by Al^3+^, AOA, malate and inhibition by vigabatrin for TaALMT1 expressed in *Xenopus laevis* oocytes and changes in [GABA]_i_. **(A)** TaALMT1 expressing oocytes under TEVC showing inward current activation (at - 80mV) in response to 100 µM Al^3+^ and 1 mM AOA addition to the bath (pH 4.5) (continuous perfusion). Rates of activation of the inward current by Al^3+^ and AOA are similar. **(B)** Current responses of TaALMT1^F213C^ mutant expressing oocytes under TEVC as in (A) Rates of activation of the inward current by Al^3+^ and AOA are similar. Currents were consistently larger for TaALMT1^F213C^ and less cRNA was injected for these oocytes (16 ng compared to 32 ng in A) **(C)** Vigabatrin (100 µM) rapidly inhibited malate-activated inward currents at pH 7.5. **(D)** Water injected control oocytes did not show a response to Al^3+^, AOA or vigabatrin. **(E)** Al^3+^ (100 µM) treatment of only 10 min caused a significant reduction in [GABA]_i_ (volume basis) only in TaALMT1 expressing oocytes. **(F)** AOA (1 mM) treatment (10 min) causes a significant reduction in [GABA]_i_ (volume basis) only in TaALMT1 expressing oocytes. **(G)** Only TaALMT1 expressing oocytes show a significant increase in [GABA]_i_ after 10 min treatment with vigabatrin (100 µM) in the presence of Na_2_SO_4_ at pH 7.5. (See Ramesh et al., (2015) for current activation by Na_2_SO_4_ at pH 7.5). Different letter indicates significance (P<0.05) (Tukey post test, One way ANOVA). For E, F and G, data are *n*=4-5 replicates (batches of 4 oocytes).

### TaALMT1 and other ALMTs also facilitate GABA influx

To test if TaALMT1 also facilitated GABA influx we expressed *TaALMT1* and *TaALMT1*^F213C^ in *X. laevis* oocytes and tested their ability to influx [^14^C] GABA. The GABA transporter AtGAT1 from *Arabidopsis* was used as a positive control in these uptake experiments (Figure 6). At pH 4.5, *TaALMT1* expressing oocytes showed significantly higher GABA uptake from an external concentration of 1 mM compared to both control and mutant *TaALMT1*^*F213C*^ expressing oocytes and similar to that facilitated by AtGAT1 (Figure 6A). At pH 7.5 *TaALMT1*-expressing oocytes (without a transactivation anion) did not show significant GABA influx compared to control oocytes, with the rate being significantly lower than *AtGAT1*-expressing oocytes (Figure 6B). *AtGAT1* expressing oocytes had significantly reduced influx at pH 7.5 compared to that at pH 4.5, consistent with the hypothesis that this transporter uses the proton motive force to drive transport (Meyer et al., 2006). Activation of TaALMT1 at pH 7.5 with 10 mM Na_2_SO_4_ increased GABA uptake significantly into the TaALMT1 expressing oocytes when compared to either the controls or *TaALMT1*^*F213C*^ expressing oocytes (Figure 6B).

**Figure 6.**
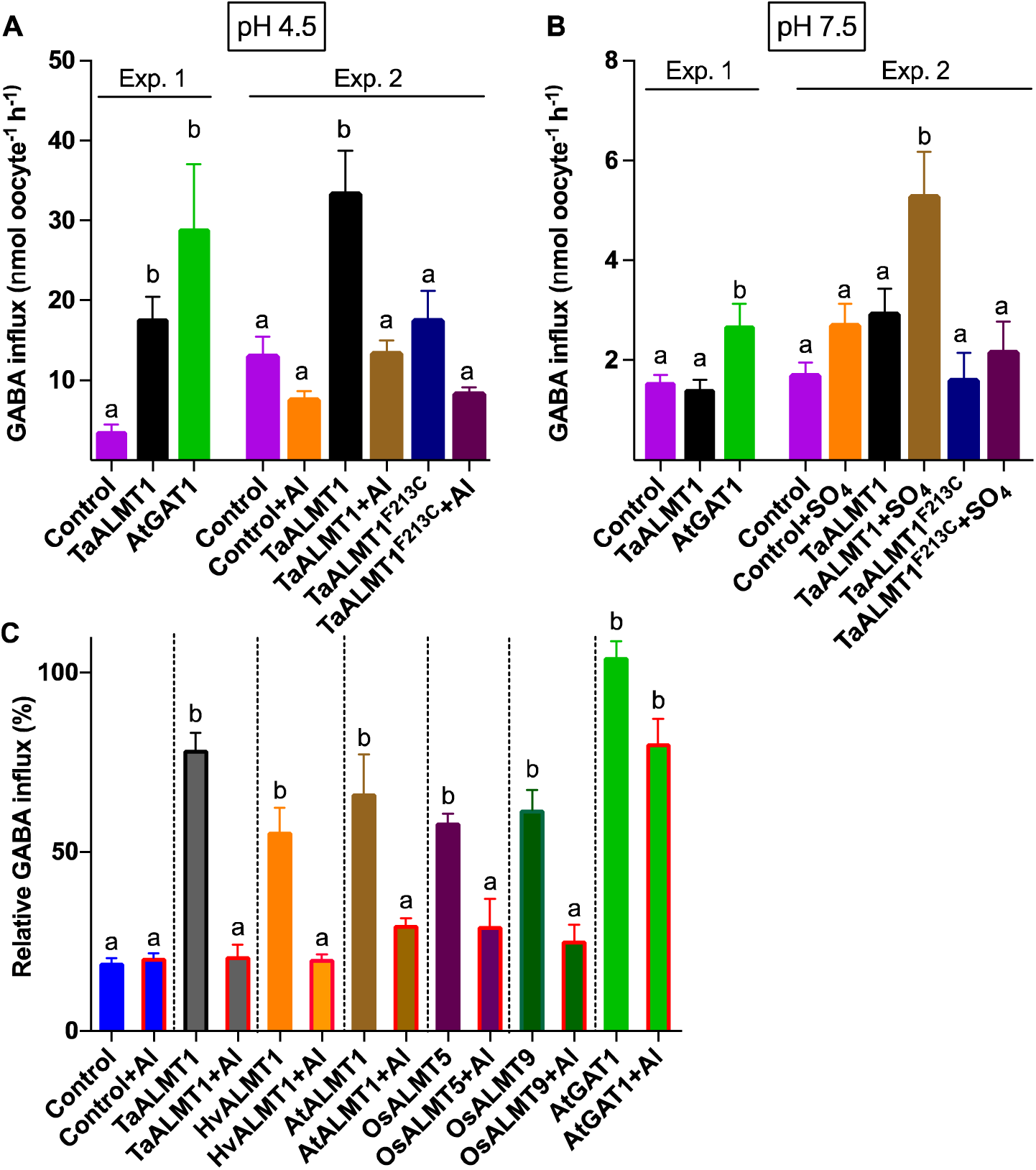
ALMTs but not TaALMT1^F213C^ facilitate pH-dependent GABA influx from 1 mM external GABA when expressed in *Xenopus laevis* oocytes and are blocked by 100 µM Al^3+^. **(A)** Comparison of ^14^C GABA uptake into *X. laevis* oocytes expressing TaALMT1 and AtGAT1 high affinity GABA transporter at pH 4.5 (Exp. 1). Exp. 2: TaALMT1 expressing oocytes showed significantly higher influx of GABA when compared with the mutant TaALMT1^F213C^ or control (water injected). There was a significant reduction in GABA uptake on addition of 100 µM Al^3+^ for oocytes expressing TaALMT1. **(B)** Comparison of ^14^C GABA uptake for TaALMT1 and AtGAT1 at pH 7.5 (Exp. 1). Exp. 2: TaALMT1 expressing oocytes showed significantly higher influx of GABA when activated by 10 mM Na_2_SO_4_ compared with the TaALMT1^F213C^ or control (water injected) oocytes. **(C)** GABA uptake was significantly higher for oocytes expressing ALMTs from wheat (TaALMT1), barley (HvALMT1), Arabidopsis (AtALMT1), rice (OsALMT5 and 9) and Arabidopsis GABA transporter (AtGAT1) at pH 4.5. Fluxes were normalised to the median of AtGAT1. Three different batches of oocytes (n=7-10). Different letters indicate significant differences between treatments at P< 0.05 Tukey post test from one-way ANOVA.

Wheat ALMT1 is the founding member of the ALMT family (Sasaki T, 2004). A large number of these genes are present in *Arabidopsis*, barley, and rice as well as other plants. Not all ALMTs are activated by Al^3+^ but can be activated by millimolar concentrations of external anions (Ramesh et al., 2015) and play diverse roles in plant physiology (Roelfsema et al., 2012; Sharma et al., 2016). Thus we tested ALMTs from *Arabidopsis*, barley and rice that had previously been shown to exhibit GABA-inhibited malate currents, for their ability to transport GABA into *X. laevis* oocytes. All the ALMTs tested (HvALMT1, AtALMT1, OsALMT5, OsALMT9) facilitated transport of GABA into the respective cRNA injected oocytes (Figure 6C). In all cases 100 µM Al^3+^ reduced this uptake to control levels. The presence of Al^3+^, however, did not affect the uptake of GABA by AtGAT1. These results demonstrate that transport of GABA is a general feature of the ALMT family and that extracellular Al^3+^ is a common inhibitor for GABA uptake of this transport across members of the family despite differences in Al^3+^-activated malate currents – of those tested only AtALMT1 and TaALMT1 have malate efflux activated by Al^3+^.

### TaALMT1 complements growth of a yeast mutant deficient in the transport of GABA

To further explore GABA transport by TaALMT1 we tested if GABA could be used as a nitrogen source for yeast growth, where TaALMT1 and site directed mutant TaALMT1^F213C^ was transformed into a yeast triple mutant strain 22754d (MATα ura3–1, gap1–1, put4–1, uga4–1) (Figure 7). This yeast strain carries mutations in general amino acid permease (gap1), proline (put4) and GABA (uga4) and is unable to grow on proline, citrulline or GABA as the sole nitrogen source; it was used to characterise high affinity GABA transport (Meyer et al., 2006). The efflux of malate was observed to be higher in each of the transformants expressing *TaALMT1* and the mutant (Figure 7A), consistent with TaALMT1 and its mutant being located to the plasma membrane and able to efflux malate, as previously shown in *X. laevis* and tobacco BY2 cells (Ramesh et al., 2015). All the yeast strains were capable of growth on selective drop out medium (SC-ura) supplemented with 2% glucose or galactose as carbon source and ammonium sulphate as nitrogen source (Supplemental Figure 9A,B). When the yeast strains were starved of nitrogen by growth in nitrogen free medium and transferred to medium with no GABA, there were no significant differences in growth of the yeast strains (Supplemental Figure 9C). However in medium supplemented with 1 mM GABA as the sole nitrogen source, yeast cells expressing TaALMT1 showed significantly higher relative growth rate compared to control (empty vector) or mutant TaALMT1^F213C^ (Figure 7B, Supplemental Figure 9E). On medium supplemented with GABA at a concentration of 20 or 37.83 mM, which corresponds to SC medium with GABA at the concentration equivalent to that of ammonium sulphate, all the yeast strains showed similar growth (Supplemental Figure 9D) indicating that the stimulation of growth by 1 mM GABA was already saturating. Therefore we examined the GABA dose-response of yeast growth (Figure 7C). The apparent K_d_ for growth stimulation by GABA was 0.56 mM indicating at least moderate affinity for GABA transport by TaALMT1. This growth stimulation was completely inhibited by the addition of 2 mM malate to the medium for TaALMT1 (Figure 7D).

**Figure 7.**
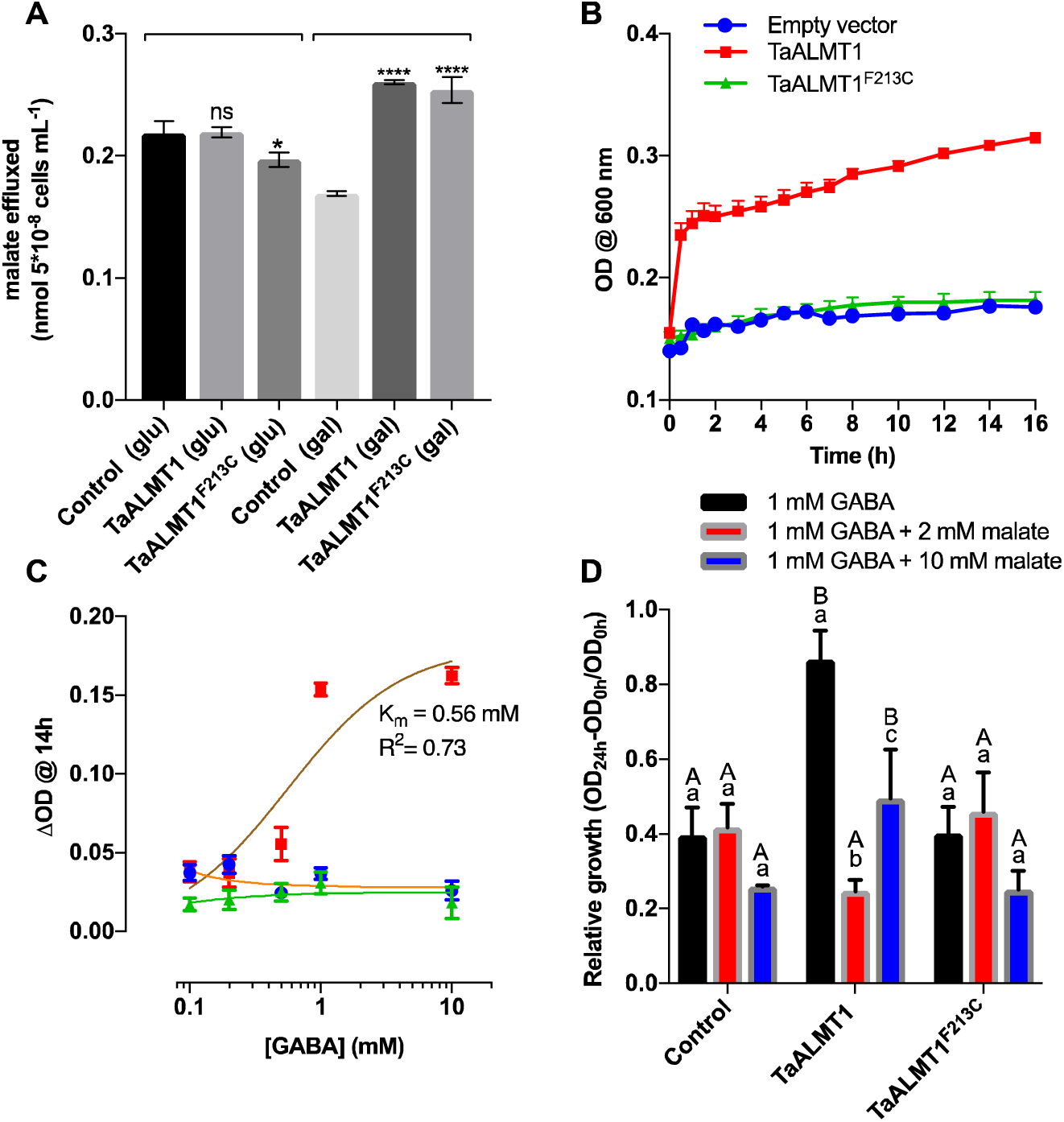
TaALMT1, but not TaALMT1^F213C^ mutant, can complement a yeast mutant strain deficient in GABA transport for growth on GABA as sole source of nitrogen. **(A)** Malate efflux over 16h from yeast mutant strain 22574d expressing TaALMT1 or F213C mutant and empty vector control grown on glucose and galactose. With galactose TaALMT1 and F213C showed increased malate efflux consistent with expression of TaALMT1 and F213C in the plasma membrane. Significance is indicated for within glucose (glu) or galactose (gal). **(B)** Growth of yeast mutant strain 22574d expressing TaALMT1 or site directed mutant F213C in medium with 1 mM GABA and galactose for 16 h. **(C)** GABA concentration dependence of yeast growth. The fitted curve is the hyperbolic Y=B_max_*[GABA]/(K_d_ + [GABA]) with best fit K_d_ of 0.56 mM. Error bars are SEM (n=6). **(D)** Comparison of the effect of 1 mM GABA on growth for empty vector controls, TaALMT1 and F213C expressing 22574d yeast strain. Presence of GABA stimulates growth only of TaALMT1 expressing yeast, which is inhibited by external malate. Small letters comparison within genotype between treatments; large letters comparison between genotypes within treatment (P<0.05) (n=3).

## DISCUSSION

### GABA transport by TALMT1 is demonstrated across a wide range of expression systems

Anion transport via ALMTs is an important and well accepted mechanism for signalling (Bouche et al., 2003a; Roberts, 2007; Pineros et al., 2008; Dreyer et al., 2013; Ligaba et al., 2013; Ramesh et al., 2015), stomatal pore regulation (Meyer et al., 2010; Roelfsema et al., 2012; De Angeli et al., 2013; Kollist et al., 2014; Palmer et al., 2016), phosphorous nutrition (Liang et al., 2013; Balzergue et al., 2017) and for providing tolerance to Al^3+^ at low pH (Delhaize and Ryan, 1995; Sasaki T, 2004; Zhang et al., 2008; Ryan et al., 2011). We have demonstrated that TaALMT1 is able to facilitate GABA efflux and influx at very high rates. This accounts for the negative linear relationship observed between malate efflux and endogenous GABA concentrations in cells expressing TaALMT1. We have shown GABA transport by TaALMT1 using near isogenic lines of wheat that differ in the expression of *TaALMT1*, transgenic barley expressing *TaALMT1*, tobacco BY2 cells expressing *TaALMT1*, *X. laevis* oocytes expressing *TaALMT1*, and complementation by TaALMT1 of a yeast mutant deficient in GABA transport. The yeast mutant 22574d that we used to characterise GABA transport and nitrogen utilisation in yeast cells (Grenson et al., 1970; Breitkreuz et al., 1999; Meyer et al., 2006) will likely be a very useful system in which to characterise other ALMTs for interactions with GABA and to further explore the pharmacology of the transporter.

### GABA transport may be a general feature of ALMTs

Uptake studies with *X. laevis* oocytes showed that GABA uptake via ALMTs is not unique to TaALMT1, but ALMTs from barley, rice and *Arabidopsis* also transported significantly more GABA into the cells than controls. Interestingly, the addition of Al^3+^ reduced the uptake in each case. We applied Al^3+^ since this activates TaALMT1 to efflux both malate and GABA at low pH. The block of GABA influx by Al^3+^ indicates that influx is mutually exclusive of efflux. In contrast the uptake of GABA by AtGAT1 was not reduced in response to Al^3+^. It is interesting to note that there is evidence for interactions of Al^3+^ with GABA_A_ receptors (Trombley, 1998) and GABA transporters in animals (Albrecht and Norenberg, 1991; Cordeiro et al., 2003).

### GABA the regulator?

Initially we investigated the role of GABA in regulating TaALMT1 based on the negative linear relationship between malate efflux and [GABA]_i_ observed in an earlier study (Figure 1 in Ramesh et al., 2015) that was hypothesised to be linked to the control of ALMT facilitated anion efflux. It was shown that there is a motif in ALMTs that is likely to be involved in GABA binding. This motif with an important phenylalanine residue (213 in TaALMT1) has similarities to a region in GABA_A_ receptors (Smith and Olsen, 1995; Boileau et al., 1999). Furthermore, some important pharmacological agents used to probe GABA receptors in animal cells also appear to work on ALMTs (e.g. muscimol and bicuculline). AtGAT1, the high affinity Arabidopsis GABA transporter used as a positive control for GABA transport in the current work, has a region of similarity with this motif (Ramesh et al., 2016).

GABA inhibits malate efflux and inward membrane currents through ALMTs when presented to the external (apoplast) side with high affinity (IC50 1 to 7 µM) (Ramesh et al., 2015). This posed the question as to how GABA is exported from the cell to the apoplast to regulate TaALMT1 if external GABA regulates ALMTs, at least for those ALMTs expressed on the plasma membrane. As far as we are aware no transport system has been identified until now that is able to account for GABA efflux across the plasma membrane into the apoplast (Shelp and Zarei, 2017). This is despite numerous examples of relatively high extracellular concentrations of GABA being measured (Chung I, 1992; Snedden et al., 1992; Crawford et al., 1994) and particularly the case for wheat roots where GABA is by far the most exported amino acid from roots (Warren, 2015).

### The location of the F213 that is critical for GABA transport

We note that there is some controversy regarding the membrane topology of ALMT proteins (Sharma et al., 2016) and this is important in the context of the location of F213 within the GABA motif of TaALMT1, and the rapidity of effect of externally applied GABA (Ramesh et al., 2015). Most recent algorithms (e.g. MEMSAT-SVM, TOPCONS, (Nugent and Jones, 2012; Tsirigos et al., 2015)) predict 6-7 transmembrane domains for the more hydrophobic N-terminus half of the protein, and with the hydrophilic C-terminus located in the cytosol. This is consistent with the model proposed by (Zhang et al., 2013) in their study of the putative pore forming domains of the vacuolar anion channel, AtALMT9. A similar model was also presented by (Ramesh et al., 2016) (reproduced in Figure 8) computed from evolutionary sequence variation over 3688 alignments using the EvFold web portal (Marks et al., 2012). Noteably the GABA motif previously characterised is located towards the end of TMD 6 (or 7) just before the long C terminus, which in most predictions is oriented toward the cytoplasm. It was suggested that this was oriented towards the apoplast to account for the rapid effect of GABA (Ramesh et al., 2015) and corresponding with immunocytochemical evidence for the C terminus to be located on the apoplast side (Motoda et al., 2007). Another model formulated from extensive sequence alignments across the family and secondary structure predictions also indicate that the GABA motif may be oriented toward the apoplast, but with a further two TMDs within the C-terminus (Dreyer et al., 2012). If the GABA motif (beginning at F213 in TaALMT1) does orient towards the cytosolic side, the transport of GABA that we demonstrate here may reconcile the rapid action of GABA on malate currents, which presumably also applies to some GABA analogs that block depending on F213, such as muscimol and vigabatrin. Clearly a crystal structure will resolve this issue.

**Figure 8.**
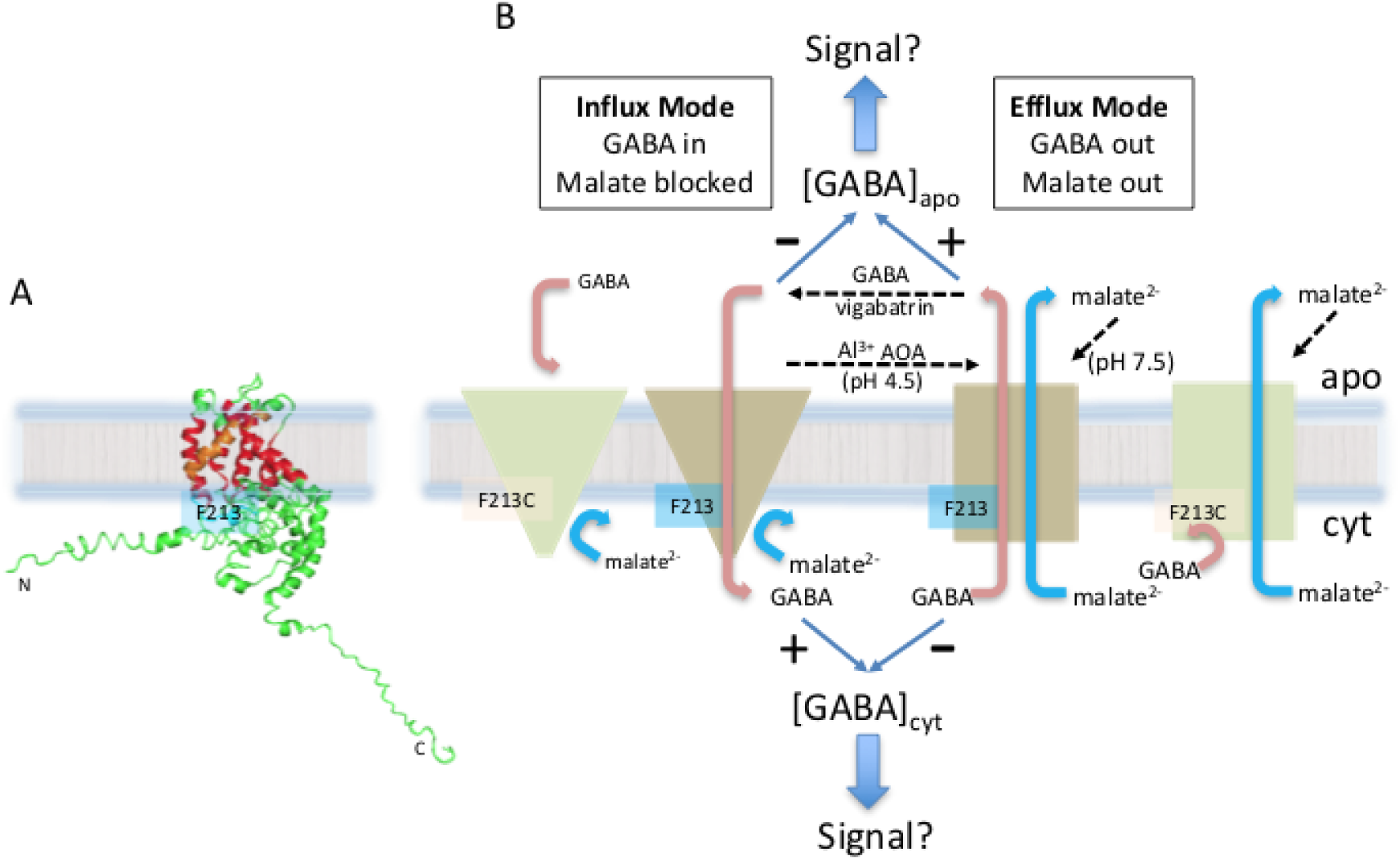
Summary of GABA and malate transport interactions through TaALMT1 and proposed sidedness of F213. **(A)** The majority of prediction algorithms (see text) indicate that TaALMT1 consists of 6 transmembrane helices (red, orange) with F213 oriented towards the cytosol at the end of the 6^th^ TM (orange, GABA motif=magenta). The N and C termini are predicted also to be on the cytosolic side. The model structure of TaALMT1 was computed from evolutionary sequence variation using the EvFold web portal (http://evfold.org/evfold-web/evfold.d) (Marks et al., 2012; Marks, et al., 2011, Ramesh et al., 2016). **(B)** Summary scheme of transport interactions and gating modes of TaALMT1. Left hand modes occur when GABA reaches a higher concentration on the outside blocking malate transport both outwards and inwards (bassed on TEVC), but GABA transport can proceed inwards. Right hand side modes occur when TaALMT1 is activated by Al^3+^ or AOA at pH 4.5 or the presence of anions (malate^2−^, SO_4_^2−^) on the outside at high pH (>7.5). The F213C mutant is shown on the outer left and right. Malate currents are activated by Al^3+^ and AOA, but are unresponsive to GABA and it does not transport GABA inwards.

### Inhibitors that target GABA metabolism and catabolism directly interact with TaALMT1

Manipulating [GABA]_i_ in cells and studying the effects of this perturbation on anion transport may provide information on the role of GABA in ALMT regulation for signalling and growth. Arabidopsis mutants deficient in GABA metabolism and catabolism have been previously used to demonstrate the role of altered [GABA]_i_ in directing pollen tube growth, guard cell closure under drought and systemic signalling upon wounding (Palanivelu et al., 2003; Mekonnen et al., 2016; Scholz et al., 2017). In this study, we used inhibitors assumed to disturb key steps in the GABA shunt pathway because of the rapidity of their effects, albeit compromised by possible lack of target specificity. Inhibition of GAD enzyme activity by AOA has been indicated for isolated mesophyll cells of Asparagus (Snedden et al., 1992). Vigabatrin on the other hand is a known inhibitor of GABA-transaminase in animal cells that catalyses the breakdown of GABA into succinate (Ben-Menachem, 2011). Both of these agents superficially appeared to have the desired affects in that AOA reduced [GABA]_i_ and activated malate efflux when *TaALMT1* was expressed, while vigabatrin increased [GABA]_i_ and inhibited malate efflux.

Apart from the expected associations with the inhibitors described above, we also observed that activation of the TaALMT1 mediated malate efflux by external Al^3+^ at low pH and by SO_4_^2−^ at pH 7.5 also reduced [GABA]_i_. This could be explained based on the prevailing view that both these treatments activate TaALMT1 directly; [GABA]_i_ may be depleted in order to support malate synthesis through the GABA shunt and the TCA cycle, since endogenous malate concentrations remain stable when malate efflux is stimulated by Al^3+^ in wheat root apices (Delhaize et al., 1993). If GABA is required for malate synthesis and the demand from increased malate efflux thought TaALMT1 results in depletion [GABA]_i_ we should expect to see a reduction in [GABA]_i_ when the TaALMT1^F213C^ mutant channel is activated. This was not observed in BY2 cells, where [GABA]_i_ remained at similar levels to controls when malate efflux was activated resulting in no relationship between [GABA]_i_ and malate efflux (Figure 3).

A direct effect of Al^3+^ and anions on the TaALMT1 protein and some other ALMTs has been demonstrated by the rapid activation observed through continuous current recording of voltage-clamped *X. laevis* oocytes (Sasaki T, 2004; Hoekenga et al., 2006; Kobayashi et al., 2007; Kovermann P, 2007.; Pineros et al., 2008; Meyer et al., 2011; De Angeli et al., 2013; Ramesh et al., 2015). Similarly GABA and muscimol inhibition of the currents was as rapid as could be expected for solution change kinetics (Ramesh et al., 2015). We have shown with *X. laevis* oocytes that AOA at pH 4.5 will rapidly activate inward current at negative membrane potential consistent with malate anion efflux accounting for the inward current. The activation occurs with the same velocity (initial current rise) as that induced by Al^3+^. Within 10 minutes [GABA]_i_ in the oocytes was significantly reduced by AOA, exactly as is observed when TaALMT1 was activated by Al^3+^. The fact that neither of these treatments changed [GABA]_i_ in control water injected oocytes indicates that the change in [GABA]_i_ was due to activation of TaALMT1. Also we observed rapid increase in inward currents in response to AOA in *X. laevis* oocytes and malate efflux from BY2 cells expressing the mutant TaALMT1^F213C^ that is not responsive to external GABA under low pH. Thus, AOA is interacting with site/s on TaALMT1 independently from the GABA binding motif.

Vigabatrin also inhibits the TaALMT1 inward current carried by anions rather rapidly and not consistent with increasing [GABA]_i_; though this inhibitor did increase [GABA]_i_ as expected in both wheat root tips and tobacco BY2 cells not expressing TaALMT1. Thus we conclude that both AOA and vigabatrin can not only alter [GABA]_i_ by their known target enzymes, but they also directly target TaALMT1 to change GABA efflux. This also illustrates the dangers of using inhibitors from animal systems to interpret functions of transporters and enzymes in plants. The negative correlations observed between malate efflux and [GABA]_i_ in both wheat roots and tobacco BY2 cells is mainly indicative of altered GABA efflux through TaALMT1. Furthermore, higher expression of *TaALMT1* appeared to elevate [GABA]_i_ independently of any treatments as was initially observed when comparing ET8 with ES8 NILs (Figure 1 in Ramesh et al., 2015) and as was observed here with tobacco BY2 cells expressing *TaALMT1* (Supplemental Figure 3) particularly evident at low pH. This is potentially due to increased accumulation of GABA via influx through TaALMT1 when TaALMT1 is not activated for “efflux mode” as was demonstrated by its capacity to influx GABA at low pH when malate efflux was not activated.

It may not be surprising that both AOA and vigabatrin interact with TaALMT1. In the case of AOA it inhibits GAD and GABA–T via interaction with the pyridoxal phosphate cofactor binding site (Wallach, 1961; John and Charteris, 1978; Löscher et al., 1989; Miller et al., 1991). It is interesting that AOA strongly activates TaALMT1, so far the only known external organic compound besides transported anions that has this effect. AOA is a structural analog of GABA that is one atom shorter and is also a zwitterion that has no net change at neutral pH. The possibility that ALMTs also have a pyridoxal phosphate binding site needs to be explored, though there is no clear pyridoxal phosphate (Vitamin B6) binding motif (InterPro IPR021115, Prosite entry Ps00392) evident in TaALMT1. GABA transaminase is also a pyridoxal phosphate dependent enzyme and is reported to be inhibited by AOA (Storici et al., 2004). Vigabatrin has a similar structure to GABA and therefore it may inhibit TaALMT1 as does the GABA analog muscimol. We conclude that the effect of vigabatrin and AOA on [GABA]_i_ is via interactions with TaALMT1, though in the case of vigabatrin it is also inhibiting GABA-T since [GABA]_i_ was increased in the absence of TaALMT1 expression.

### Potential roles for GABA transport via ALMTs

Until now the identity of a GABA efflux transporter across the plasma membrane was not known. Our studies show that TaALMT1 in addition to mediating efflux of organic anions also mediates GABA efflux from cells when activated. Its transport capacity for GABA is very high as indicated by its activation being able to significantly reduce [GABA]_i_ in root tips and other cells expressing TaALMT1. In fact, we have shown higher capacity for GABA efflux relative to malate efflux on a molar basis. This is quite novel as changes in [GABA]_i_ are likely to have broad effects on carbon metabolism and signalling. We have shown that GABA does not significantly complex Al^3+^ therefore we can exclude the hypothesis that GABA efflux with malate may provide additional protection against Al^3+^. Although it is not clear why high GABA efflux may be an advantage when TaALMT1 is activated by Al^3+^ at low pH, or activated at high pH, we may speculate along the following lines as follows.

Firstly GABA may act as an extracellular pH buffer. GABA addition to solutions at both low and high pH tends to bring the pH back towards neutrality. Thus at low pH where GABA synthesis by GAD acts as a cytoplasmic pH stat (Snedden et al., 1992; Crawford et al., 1994; Shelp et al., 1999; Snedden and Fromm, 1999), its efflux to the external medium will also tend to increase the external pH. It has been reported that exogenously supplied GABA (10

µM) significantly improved root growth of barley seedlings at pH 4.5 and when exposed to 20 µM Al^3+^ at pH 5 (Song et al., 2010). This was considered to be via alleviation of oxidative damage.

Secondly AtALMT1 has been implicated as the key regulator in the attraction of the beneficial *Bacillus subtilis* strain FB17 to the Arabidopsis root (Lakshmanan et al., 2013), in addition to its role in Al^3+^ tolerance. Pseudomonads are known for their specific GABA receptors and positive chemotactic response to GABA (Reyes-Darias et al., 2015) and *Bacillus subtilus* has positive chemotaxis to Arabidopsis roots with chemoreceptors that recognise a range of amino acids.

Thirdly GABA transported by TaALMT1 also provides feedback regulation on the co-transport of organic anions. As GABA builds up in the apoplast this will tend to reduce malate efflux. Apoplastic GABA may also signal to adjacent cells via its effect on TaALMT1. Malate has the opposite effect since it activates TaALMT1 from the *trans* side of TaAMT1 particularly at higher pH. Thus the two transported substrates have opposite effects on the *trans* (apoplastic) side of the transporter (Figure 8). ALMTs in general have rather complex regulation via anions, for example the regulation of Cl^−^ efflux for stomatal closure in the date palm by external NO_3_ ^−^ (Müller et al., 2017), and the activation of AtALMT9 on the tonoplast membrane by cytosolic malate to efflux Cl^−^ into the vacuole of guard cells during stomatal opening (De Angeli et al., 2013).

### The transport mechanism for GABA and its interaction with malate anions

Considering the transport of GABA via TaALMT1, it is necessary to account for the likely ionic charge on GABA. GABA is a zwitterion at neutral pH and is thus uncharged in the cytosol. Only at very high pH (10.43) or at low pH (4.23) does either a negative or positive charge occur respectively. At pH 4.5 we calculate that there would be 35% of GABA in the external media that has a net positive charge due to a proportion of molecules not deprotonated at the carboxyl end while the amino terminal will remain positively charged.

Efflux from the cytoplasm (slightly alkaline pH) would not be detectable as an electrical current, but influx at pH 4.5 could be detected as an inward current if the cationic form of GABA is transported, particularly if there is a high affinity of transport as suggested by the IC_50_ of block by GABA of the anion current, and from the high influx measured relative to that of AtGAT1 at pH 4.5. It is also possible that GABA fluxes may be coupled to the movement of protons in the same direction similar to GABA transport via AtGAT1 (Meyer et al., 2006), which would be detectable as an electrogenic current. GABA influx into *X. laevis* oocytes was substantially higher at pH 4.5 than pH 7.5 (10-fold), and the flux at pH 7.5 was similar to that of control water injected oocytes. However, GABA influx was activated at pH 7.5 by the addition of Na_2_SO_4_ to the bathing medium, which also stimulates malate efflux.

We have shown that for all the ALMTs that tested positive for GABA influx, external GABA also inhibited the anion efflux current at pH 7.5 at micromolar concentrations (Ramesh et al., 2015). This may indicate some coupling between anion efflux and GABA transport. GABA on the *trans* side for anions (normally the external side of the plasma membrane) inhibits anion efflux but results in GABA uptake. Malate on the *trans* side also inhibits GABA influx. However, GABA on the inside of the cell clearly allows efflux of anions and GABA together, since in the heterologous expression systems used here and in the wheat NIL lines both GABA and malate efflux occur simultaneously. In the case of *X. laevis* oocyte the [GABA]_i_ is well above the concentrations that blocks malate efflux from the *trans* side. The F213C mutation in TaALMT1 greatly reduces the external GABA sensitivity of anion efflux currents (Ramesh et al., 2015). We have shown here that this mutation also effectively abolishes GABA influx and GABA efflux via TaALMT1, but still allows activation by external anions and external Al^3+^, further confirming the importance of this site in GABA interactions. The F213C mutation also shows extremely high malate efflux when expressed in *X. laevis* oocytes compared to the unmutated TaALMT1, suggesting that “co-transport” of malate and GABA has been uncoupled. A summary of the observed interactions between GABA and malate transport via TaALMT1 is presented in Figure 8.

To fully understand the transport mechanism in ALMTs more detailed kinetic studies will be required where GABA and malate concentrations are varied on both *cis* and *trans* faces, most probably best achieved via patch clamp studies. Our work reported here indicates that ALMTs are more complicated than a relatively simple ligand gated anion channel, and we suggest that at least some (e.g. TaALMT1) could be considered as GABA transporters with anion channel activity. Why this has not be revealed from the many previous studies is probably related to several factors including the focus upon Al^3+^ tolerance resulting from the malate transport through TaALMT1 and the probable lack of or low electrogenic activity associated with GABA transport.

The dual function of TaALMT1 and other ALMTs is analogous to some transporters that display channel activity such as the excitatory amino acid transporters (EAATs) from animals that function as both glutamate transporters and chloride channels (Cater et al., 2016). There is also a precedent for GABA transport in that the mammalian GAT1 transporter displays sodium channel activity (Risso et al., 1996). In the context of the biological link between GABA and anion transport and particularly that of malate, the regulation of efflux of both substrates is titrated finely by apoplastic concentrations of both substrates making them uniquely positioned to provide intercellular and intracellular communication of metabolic status (Figure 8).

## METHODS

### Chemicals

All chemicals were purchased from Sigma. ^14^C GABA was obtained from American Radiolabelled Chemicals, Inc, USA.

### cRNA synthesis

Plasmid DNA from ALMT and site-directed mutants (Nugent and Jones, 2012; Ramesh et al., 2015) was extracted using the Mini Prep kit from Sigma, and 1 μg of plasmid DNA was linearized with the restriction enzyme Nhe1. Capped complementary RNA (cRNA) was synthesized using the MESSAGE Mmachine T7 Kit (Ambion) as per the manufacturer’s instructions.

### Voltage-clamp electrophysiology

Electrophysiology was performed on *X. laevis* oocytes 2 days post injection with water/cRNA (Preuss et al., 2010; Ramesh et al., 2015). Oocytes were injected with 46 nl of RNase-free water using a micro-injector (Nanoject II, automatic nanolitre injector, Drummond Scientific) ± 16–32 ng cRNA. Sodium malate (10 mM, pH 7.5) was injected into oocytes 1 h before measurement. Aluminium activation was carried out in ND88 solution at pH 4.5 (Sasaki T, 2004; Hoekenga et al., 2006) ± aluminium chloride (AlCl_3_-100 µM) and vigabatrin (100 µM). Basal external solutions for anion activation contained 0.5 mM CaCl_2_ (pH 4.5) or 0.7 mM CaCl_2_ (pH 7.5) and mannitol to 220 mOsm kg^−1^, ± 10 mM sodium sulphate or 10 mM malate and amino oxyacetate (AOA-1 mM) buffered with 5 mM BTP/MES from pH 4.5 to 7.5. In all *X. laevis* oocyte experiments, solutions were applied to gene-injected oocytes in the same order as controls (water injected). Randomly selected oocytes were alternated between control and gene injected to limit any bias caused by time-dependent changes after gene injection or malate injection. The University of Adelaide Animal Ethics Committee approved the *Xenopus laevis* oocyte experiments; project number S-2014-192.

### Root assays

NILs of wheat ET8 and ES8 (Sasaki T, 2004) and barley transgenic line overexpressing TaALMT1 - a gift from Dr. Peter Ryan, CSIRO, Canberra (Delhaize et al., 2004) were surface sterilized, and 3-day-old seedlings were placed in a microcentrifuge tube with roots immersed for 22 h in 3 mM CaCl_2_, 5 mM MES/BTP to pH 4.5–7.5 ± treatments. For root flux assays, experiments were carried out wherein the identity of the treatment solutions was unknown to the person performing the experiments to remove any bias. Malate concentrations were measured on an OMEGA plate-reading spectrophotometer (BMG) following the K-LMALR/K-LMALL assay11 (Megazyme). One hundred microlitres of the treatment solution collected from roots, was added to a mastermix containing the various components of the K-LMALR/K-LMALL assay11 (Megazyme) kit as per the manufacturer’s instructions. The change in absorbance at 340 nm was used to calculate the concentration of malate in the samples.

### Endogenous GABA concentrations and GABA efflux

GABA concentrations were measured, on the OMEGA plate-reading spectrophotometer, following the GABase enzyme assay (Zhang and Bown, 1997). Briefly, 5 mm of root tips were excised and snap frozen in liquid nitrogen after seedlings were subjected to treatment solutions for 22 h unless otherwise stated. The root tips (3 per seedling / replicate) were ground in liquid nitrogen and known weight (10-15 mg) was added to methanol and incubated at 25 °C for 10 min. The samples were vacuum dried, resuspended in 70 mM LaCl_3_, pelleted at 500 x g in a desktop microcentrifuge and precipitated with 1M KOH. These samples were re-centrifuged at 500 x g and 45.2 μl of supernatant was assayed for GABA concentrations using the GABase enzyme from Sigma as per the manufacturer’s instructions. For oocyte GABA measurements, *Xenopus* oocytes injected with TaALMT1 cRNA or water (controls) were imaged in groups of 4-5 using a stereo zoom microscope (SMZ800) with a Nikon (cDSS230) camera 48h post-injection. The oocytes were incubated in treatment solutions indicated in the figure legends for 10 min. After 10 min, treatment solution was removed and oocytes snap frozen in liquid nitrogen and stored at −80 °C until further use. Oocytes were extracted with same protocol used for root tips and assayed as mentioned above. Samples from root flux assays or tobacco BY2 suspension cell assays were used to measure external GABA using the GABase enzyme as described above.

GABA concentrations were also analysed with UPLC. As described above the same GABA extracts from root tips, oocytes, and tobacco BY2 cells were centrifuged at 16,000 x g in a table top centrifuge for 10 minutes, and the supernatant was filtered with a 0.2 μm syringe filter. For GABA efflux, 1 mL of efflux solution was concentrated by drying down and reconstituted with 200-250 μL of water, followed by the filtering step as mentioned above. GABA in the samples was derivatized with the AccQ•Tag Ultra Derivatization Kit (Waters, USA) according to the manufacturer’s protocol. Chromatographic analysis of GABA was performed on an ACQUITY UPLC System (Waters, USA) using a Cortecs^®^ UPLC C_18_ column (1.6 μm, 2.1 × 100 mm). The gradient protocol for amino acids analysis was used to measure GABA with mobile solvents AccQ•Tag Ultra Eluents A and B (Waters, USA). Calibration range for GABA was 0 to 500 μM. The results were analysed with Empower 3 chromatography software by Waters.

### Tobacco BY2 transgenic lines

Tobacco suspension cells (*Nicotiana tabacum* L. cv. Bright Yellow 2) were transformed with TaALMT1, site directed mutants or an empty vector pTOOL37 using a slightly modified protocol (An, 1985). Briefly, fresh suspension cells in Murashige and Skoog’s (MS) medium were co-cultivated with agrobacterium strain carrying the binary vectors with genes of interest for 48 h with 20 uM acetosyringone at 25°C. The cells were washed four times in MS medium with a final wash in MS medium with carbenicillin (500 µg/ml). Cells were then plated on selective plates with MS medium, carbenicillin and hygromycin (50 µg/ml) and incubated at 25°C for 14 days. Microcalli visible on plates were transferred to fresh selective plates for further growth. The transformed calli were used to extract DNA to confirm the presence of transgenes by Polymerase Chain Reaction (PCR) and maintained by subculture every three weeks. The transgenic BY2 cell lines were used in further experiments.

### Tobacco BY2 malate efflux and GABA concentrations

Tobacco suspension cells were grown in Murashige and Skoog’s medium on a rotary shaker (~100 r.p.m.) until the logarithmic phase. Aliquots of suspension cells containing ~1 g of cells were centrifuged and gently resuspended in a basal BY2 solution (Zhang WH and SD., 2008). TaALMT1, site directed mutants-expressing or vector-control tobacco-BY2 suspension cells (0.15 g) were placed in 3 ml of 3 mM CaCl_2_, 3 mM sucrose and 5 mM MES/BTP to pH 4.5–7.5 ± treatments as shown in the figure legends, in 50-ml tubes on a rotary shaker for 22 h, unless otherwise stated. Malate fluxes were measured as stated above. Cells were harvested after treatments by centrifugation at 5000 x g in a desktop microcentrifuge and snap frozen in liquid nitrogen. Frozen cells were subjected to extraction and GABA concentrations measured as described above using GABase enzyme (Sigma)

### ^14^C GABA tracer flux experiments

*Xenopus* oocytes expressing ALMTs, site directed mutant, controls and AtGAT1 (At1g08230) were incubated in uptake buffer containing 0.5 mM CaCl_2_ (pH 4.5) or 0.7 mM CaCl_2_ (pH 7.5), buffered with 5 mM MES/BTP ± AlCl_3_ (100 µM)/sodium sulphate (10 mM), mannitol to 220 mOsm kg^−1^ and 1.018 mM GABA for 2 h. The oocytes were washed twice with ice cold uptake buffer and placed in 0.1N nitric acid before addition of scintillation fluid (4 ml) and radioactivity measured in a Beckman and Coulter multipurpose scintillation counter (LS6500). The AtGAT1 used as a positive control was a gift from Prof. Doris Rentsch, University of Bern, Switzerland.

### Yeast Experiments

*S. cerevisiae* strain 22574d (*MAT*α *ura3-1, gap1-1, put4-1, uga4-1*) (Jauniaux et al., 1987) was transformed with TaALMT1 and site directed mutants cloned into yeast transformation vector pDEST52 according to (Schiestl and Gietz, 1989) and the transformants were selected on minimal media supplemented with 2% glucose. Selected transformants were grown in Grenson’s (Grenson et al., 1970) media lacking nitrogen supplemented with 2% galactose for 16-18h before transfer to media supplemented with either 1 mM GABA as the sole nitrogen source for growth studies or different concentrations of GABA (0-1 mM) for dose response curve. The yeast strain used was a gift from Prof. Doris Rentsch, University of Bern, Switzerland.

### Ligand Complexation Method for testing Al^3+^-GABA complexation

A competitive ligand method was developed and tested wherein the ligand of interest, L (e.g. GABA), is added to a solution containing Al^3+^ and fluoride (F). Depending on the strength of Al^3+^ complexation with the ligand of interest, F will be displaced from the AlF^2+^ complex, resulting in higher free F concentration, which can be detected with a fluoride ion selective electrode (ISE).

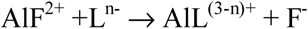

Five organic ligands were used in the measurements: citric acid, oxalic acid, malic acid, salicylic acid and GABA. Stock solutions of 0.05 and 0.005 M were made and adjusted to pH 4.5 using NaOH. A solution containing 10 mM NaCl, 60 μM Al^3+^ (as AlCl_3_), 60 μM F (as Na_2_SiF_6_) and 10 mM MES, adjusted to pH 4.5 was made. Ten ml of this solution was titrated with organic ligand, and the free F concentration was determined using a fluoride ion selective electrode (Orion 9609BNWP), allowing several minutes between each titration step until the reading stabilized. The ISE was calibrated using solutions with 10 mM NaCl, 10 mM MES (pH 4.5) and 1-100 μM F. All ISE measurements were made in Al-foil covered beakers while stirring the solution. The pH of the solution was checked at the end of each titration run and was always between 4.45 and 4.55

### Statistics

All graphs and data analysis were performed in GraphPad Prism 7 (version 7.02). All data shown are mean ± SEM.

### Accession numbers

Sequence data from this article can be found in the EMBL/GenBank data libraries under accession number(s) TaALMT1 (DQ072260); HvALMT1 (EF424084); OsALMT5 (Os04g0417000); OsALMT9 (Os10g0572100); AtALMT1 (AT1G08430); AtGAT1 (At1g08230); Actin (KC775782); Tubb4 (U76895.1); Cyclophilin (EU035525.1); GAPDH (EF592180.1).

## Supplemental Data

**Supplemental Figure 1.** GABA spike and recovery experiments with root tips of 3 day old wheat seedlings (ET8) and *Xenopus* oocytes (control and TaALMT1 injected) to test the impact of AOA, Al^3+^ and vigabatrin treatment on the enzyme assay of GABA and UPLC vs GABase mesurements.

**Supplemental Figure 2.** Expression of *TaALMT1* relative to three housekeeping genes (*GAPDH, Cyclophilin and Actin*) in seedling roots of plants treated with 100 µM AlCl_3_ pH 4.5 or basal solution for 22 hours.

**Supplemental Figure 3.** Comparison of malate efflux and endogenous GABA concentrations in BY2, empty vector (control), TaALMT1 and TaALMT1^F213C^ mutant expressing cells treated with 100 µM Al^3+^ or 1 mM AOA at pH 4.5.

**Supplemental Figure 4.** Malate efflux via TaALMT1 expressed in tobacco BY2 cells as a function of external pH and the effect of AOA.

**Supplemantal Figure 5.** Negative linear correlation between malate efflux and cell GABA concentration in Tobacco BY2 cells expressing TaALMT1 for the individual replicates for the treatments taken together from Figure 3 A,B.

**Supplemental Figure 6.** Vigabatrin, an inhibitor of GABA transaminase, inhibits malate efflux activated by 10 mM Na_2_SO_4_ while elevating endogenous GABA concentrations in BY2 cells expressing TaALMT1 at pH 7.5.

**Supplemental Figure 7.** Comparison of malate efflux and endogenous GABA concentrations in BY2, empty vector (control), TaALMT1 and TaALMT1^F213C^ mutant expressing cells treated with addition of 10 mM Na_2_SO_4_ and 100 µM vigabatrin at pH 7.5.

**Supplemental Figure 8.** Tobacco BY2 empty-vector controls show no significant correlation (P>0.05) between malate efflux and endogenous GABA concentrations at pH 4.5 when treated with Al^3+^ AOA, or at pH 7.5 when treated with Na_2_SO_4_ or vigabatrin.

**Supplemental Figure 9.** Details of complementation of yeast strain 22754d (MATα ura3–1, gap1–1, put4–1, uga4–1) expressing TaALMT, TaALMT^F213C^.

**Supplemental A.** Details of measurement of possible complexation of Al^3+^ by GABA.

## ACKNOWLEDGEMENTS

We thank Prof. Doris Renstch for providing the yeast strain and Dr. Peter Ryan for providing transgenic barley seeds. We thank Muyun Xu for cloning the cDNAs of OsALMT5 and OsALMT9. The Australian Research Council has supported this research through CE140100008 and DP130104205 awarded to S.D.T. and M.G., who is also supported by FT130100709.

## AUTHOR CONTRIBUTIONS

S.D.T supervised the research. S.A.R, S.D.T and M.G. planned and designed experiments. S.A.R performed oocyte, tobacco BY2 cells, yeast and plant experiments. M.K performed additional plant experiments. W.S. assisted in plant, oocyte and tobacco cell experiments. M.O and L.C. optimised and carried out UPLC measurements. M.M. and F.D. optimised and carried GABA-Al^3+^ binding assays. S.A.R and S.D.T analysed the data and wrote the paper with edits from M.G. All authors had intellectual input into the project and commented on the manuscript.

